# Transcriptional regulation of ACSL1 by CHREBP and NF-kappa B in macrophages during hyperglycemia and inflammation

**DOI:** 10.1101/2020.08.04.235887

**Authors:** Prashanth Thevkar-Nagesh, Shruti Rawal, Tarik Zahr, Edward A. Fisher, Michael J. Garabedian

**Affiliations:** Department of Microbiology, New York University School of Medicine, New York, NY 10016, USA; Department of Medicine, New York University School of Medicine, New York, NY 10016, USA

**Keywords:** Acyl-CoA synthetase 1 (ACSL1), transcription, CHREBP, NF-kappa B, hyperglycemia, diabetes, inflammation, macrophage

## Abstract

Acyl-CoA synthetase 1 (ACSL1) is an enzyme that converts fatty acids to acyl-CoA-derivatives for use in both lipid catabolism and lipid synthesis, including of arachidonic acid mediators that promote inflammation. ACSL1 has also been linked to the pro-atherosclerotic effects of diabetes in mice. ACSL1 expression has been reported to be upregulated in monocytes and macrophages by hyperglycemia, as well as enhanced by inflammatory stimuli, yet surprisingly little is known about the mechanisms underlying its transcriptional regulation. Here we show that increased Acsl1 mRNA expression in mouse macrophages by hyperglycemia is via transcription initiation such that nascent ACSL1 RNA and Acsl1 promoter activity are increased. We further demonstrate that the hyperglycemic-dependent induction of Acsl1 mRNA is governed by the glucose-sensing transcription factor, Carbohydrate Response Element Binding Protein (CHREBP), since the hyperglycemic upregulation of Acsl1 mRNA is lost in mouse bone marrow derived macrophages (BMDMs) from Chrebp knock out mice. In addition, we show that LPS treatment of mouse BMDMs increased Acsl1 mRNA, and this is attenuated by an NF-kappa B inhibitor that blocks p65 subunit binding to DNA. We further show that LPS treatment increased ACSL1 protein abundance and stimulated ACSL1 protein localization to membranes where it likely exerts its activity. Using an ACSL1 reporter gene containing the promoter and 1.6 Kb of upstream regulatory region, which contain multiple predicted CHREBP and NF-kappa B (RELA) binding sites conserved between the human and mouse ACSL1 gene, we found a synergistic increase of ACSL1 promoter activity when CHREBP and RELA were co-expressed. Thus, we have identified pathways controlling the expression of ACSL1 by hyperglycemia and inflammation through CHREBP and NF-kappa B.

## INTRODUCTION

Atherosclerosis is a chronic inflammatory disease characterized by infiltration and deposition of lipid laden macrophages, termed “foam cells”, in the arterial wall [1]. Circulating monocytes are recruited to the plaque by inflammatory signals, and become activated macrophages and proinflammatory, further contributing to the plaque development.

There are several factors that drive the inflammatory response of macrophages, including the enzyme ACSL1, which converts long-chain fatty acids into acyl-CoA derivatives [2, 3]. When arachidonic acid (C20:4) is taken up by macrophages it is converted by ACSL1 to arachidonoyl-CoA (20:4-CoA), which in turn is incorporated into phospholipids. This 20:4-CoA moiety can be liberated from the phospholipids by phospholipase A2 (PLA2) and made available for prostanoid production, and activation of prostaglandin-endoperoxide synthase (PTGS; *aka* COX2) to enhance inflammation [3, 4].

ACSL1 has also emerged as a mediator of the enhanced atherosclerosis associated with diabetes [3]. Diabetes increases the risk of cardiovascular disease by accelerating the progression of atherosclerosis [5]. The expression of ACSL1 mRNA has been shown to be increased in monocytes from human diabetic patients and mouse models of type-1 diabetes [4]. Moreover, this increased expression was also observed when macrophages were cultured under diabetes-relevant high glucose (25mM) compared to normal glucose (5.5mM), suggesting that the effect of hyperglycemia on ACSL1 expression is cell autonomous. Importantly, the accelerated progression of atherosclerosis under diabetes was prevented in mice lacking *Acsl1* in monocytes and macrophages [4]. Thus, ACSL1 is a key regulator of the pro-atherosclerotic effects of diabetes. Consistent with ACSL1 links to cardiovascular and metabolic disease in humans, analysis of genome wide association studies found intronic SNPs in ACSL1 associated with atherosclerosis and type-2 diabetes [6] ,

In addition to hyperglycemia, ACSL1 expression in macrophages is induced by lipopolysaccharide (LPS) and gram negative bacteria (E. Coli) [7]. Moreover, inflammatory M1 macrophages showed not only an increased in the abundance of ACSL1 protein, consistent with the increased mRNA, but greater ACSL1 protein localization to the plasma membrane relative to non-activated (M0) macrophages, suggesting that membrane location of ACSL1 is part of its proinflammatory response by aligning ACSL1’s enzymatic activity to the site of its substrates [2].

Despite the increased expression of ACSL1 in macrophages by hyperglycemic and inflammatory stimuli, little is known about the mechanisms mediating the transcriptional regulation of ACSL1, including the transcription factors controlling ACSL1 expression. Here we report the analysis of the mechanisms and transcription factors controlling ACSL1 expression. We find that transcriptional regulation of ACSL1 under hyperglycemia and inflammatory stimuli involve, respectively, CHREBP and NF-kappa B-mediated activation of the ACSL1 promoter.

## MATERIALS AND METHODS

### Cell culture

Human embryonic kidney (HEK) 293 cells (ATCC) were cultured in Dulbecco’s modified Eagle’s medium (DMEM, Corning) containing 10% fetal bovine serum (FBS) and 1% PenStrep (100 U/mL Penicillium and 100ug/mL Streptomycin) in either 4.5 g/L D-glucose or 1 g/L D-glucose + 3.5 g/L L-glucose (Sigma) to serve as an osmotic control. Cells were tested for mycoplasma and were tested negative. Cells were cultured with 5% CO_2_ at 37°C.

### Animals

Wild type mice (C57B16J) were obtained from Jackson labs. Tibias and femurs from *Mlxipl* deficient mice *(Chrebp*^***−/−***^) were kindly provided by Dr. Claudia Han from the Glass lab at UCSD. The animals were cared for in accordance with the National Institutes of Health guidelines and the NYU and UCSD Institutional Animal Care and Use Committee. Mice were euthanized by CO_2_ followed by cervical dislocation in accordance with approved guidelines for the euthanasia of animals.

### Bone marrow derived macrophages (BMDMs)

BMDMs were isolated from the tibia and femur of 6–12-week-old male C57BL6J mice. Isolated bone marrow cells were treated with red blood cell lysis buffer (Sigma) and re-suspended in differentiation medium (DMEM with 1 g/L D-glucose + 3.5 g/L L-glucose or 4.5 g/L D-glucose and L-glutamine, supplemented with 20% FBS and 10 ng/μL macrophage colony-stimulating factor (M-CSF) (PeproTech, Inc., Rocky Hill, NJ). Cells were passed through a 70 μm filter to clear any debris. Following this, cells were plated in 10cm non-tissue coated plates and allowed to differentiate for 7 days to obtain un activated (M0) macrophages. At day 7, the cells were washed in PBS, and re-plated with desired cell density in a 6-well dish and allowed to attach to the plate. Cells were treated with LPS (50ng/ml) for the indicated times, and RNA or protein was isolated. For some experiments, NF-kappa B inhibitor, CAPE (5μM) was pretreated for 4 hours before LPS treatment.

### Promoter motif prediction

Eukaryotic Promoter Database was used for predicting the putative CHREBP (*Mlxipl)* and p65 (RELA) sites on the upstream ACSL1 promoter using the mouse and human database [8].

### Luciferase assay

HEK 293 cells (24 well format) were transfected with Lipofectamine 3000 (Invitrogen) following manufacturers protocol. Cells were transfected with 250ng of vector only or ACSL1-GL or CHREBP+ ACSL1-GL or p65 (RELA) + ACSL1-GL constructs. For co-transfection experiments, p65 (RELA) (250ng) and were CHREBP (250ng) were co-expressed with ACSL1-GL and the vector only was adjusted to 750ng. At 48 hour post transfection, media was collected from the respective cells and luciferase assay was performed following manufacturer’s protocol with GL-S buffer. Luciferase activity was measured using the LMax microplate reader luminometer with an integration time of 3 sec. The ACSL1-Gaussian luciferase (GL) reporter construct and control Gaussian luciferase vectors were obtained from Genecopeia (Product ID: MPRM39476). pcDNA3 Flag-RelA was purchased from Addgene (plasmid #20012) and deposited by Stephen Smale [9]. The ChREBP expression vector was purchased from Addgene (plasmid #39235) and deposited by Isabelle Leclerc [10].

### RNA isolation, cDNA synthesis and qPCR

Total RNA was isolated using RNeasy Mini Kit (Qiagen). On-column DNase digestion step was performed during the isolation process. cDNA was synthesized from 500ng of RNA using Thermo Scientific™ Verso cDNA Synthesis Kit (AB1453B) following the manufacturer’s instructions. Quantitative real-time PCR was performed on the QuantStudio 6 Flex (Applied Biosystems) using SYBR Green Fast Master Mix (Applied Biosystems). 5ng of cDNA and 100nM primers were used for performing the qPCR reaction. Gene expression was calculated using the relative quantification (2−ΔΔCT method) or by absolute quantification using standard curve.

### Primers

ACSL1 mRNA

F 5’-GCGGAGGAGAATTCTGCATAGAGAA-3’;

R 5’-ATATCAGCACATCATCTGTGGAAG-3’

Cyclophilin A1 mRNA

F: 5’-GGCCGATGACGAGCCC-3’

R: 5’-TGTCTTTGGAACTTTGTCTGCAA-3’

ACSL1 hnRNA

F: 5’-TCACTCCTTATCACCTCTTC-3’

R: 5’-CTCCAGAGCTTTGAGGCTGATG-3’

### Immunoblotting

Cells were lysed in lysis buffer (50 mM Tris-HCl [pH 7.4], 150 mM NaCl, 0.5% sodium dodecyl sulfate [SDS], 0.5% sodium deoxycholate, 1% Triton X-100, and 1× protease inhibitor cocktail [Roche]). The total amount of protein was quantitated by using Pierce™ Rapid Gold BCA Protein Assay Kit (Thermo Scientific). Equal amounts of proteins were resolved on 10 or 15% Tris-glycine SDS-PAGE under reducing conditions and transferred onto Immobilon-P Membrane, PVDF, 0.45 μm (Millipore). Membranes were probed with rabbit anti-ACSL1 (#9189, 1:1,000; Cell Signaling), rabbit anti-CHREBP Antibody (# NB400-135, 1:500; Novus Biologicals), rabbit anti-histone H3 (1:1,000; Cell Signaling), rabbit anti-actin (1:5,000; Abcam), rabbit-anti Sodium Potassium ATPase antibody (1:5,000; ab76020, Abcam) or mouse anti-tubulin Mouse Monoclonal Antibody (HRP-66031, 1:5000, Proteintech), followed by horseradish peroxidase-conjugated anti-mouse, or anti-rabbit IgG antibody (1:5,000; Life Technologies). Protein bands were visualized by using a Clarity Western ECL Substrate (BioRad), and images were acquired on an Odyssey Fc imaging system (Li-Cor).

### Immunofluorescence

Cells were fixed in 4% methanol-free paraformaldehyde (Fisher Scientific) and permeabilized with 0.2% Triton X-100. 5% mouse or rabbit serum was used for blocking. Cells were stained with rabbit anti-CHREBP antibody (# NB400-135, 1:100; Novus Biologicals), followed by Alexa 488-conjugated goat anti-rabbit IgG secondary antibody (1:400; Invitrogen) for 1 hour at room temperature. Finally, cells were stained with the DNA-binding dye Hoechst (5 μg/ml; Invitrogen), and coverslips were mounted in mounting medium (Sigma-Aldrich). Fluorescent images were acquired by sequential scanning on a Leica SP5 Confocal Microscope confocal laser scanning microscope. Acquired images were analyzed in ImageJ.

### Statistical analysis

All statistical analyses were performed using Prism 6 (GraphPad). P values were calculated by using unpaired t tests for pairwise data comparisons, one-way analysis of variance (ANOVA), or two-way ANOVA for multiple comparisons. A P value of ≤0.05 was considered significant.

## RESULTS

### *Acsl1* expression is transcriptionally upregulated in macrophages under hyperglycemia

It has been reported that the expression of *Acsl1* is upregulated in diabetic monocytes and macrophages *in vitro, in vivo,* and in clinical samples [4, 6, 11] However, the mechanism whereby *Acsl1* is upregulated by hyperglycemia is not understood. For instance, is the increase in A*csl1* mRNA by high glucose a result of an increase in the transcriptional initiation of the gene? To interrogate the regulation of *Acsl1*, we differentiated primary mouse bone marrow derived macrophages (BMDMs) under normal glucose (NG; 5.5 mM glucose) and high glucose (HG; 25mM glucose) conditions. We found that *Acsl1* mRNA and protein were upregulated under HG compared to NG (Fig. 1A and B). To identify if this increase reflected enhanced transcription, we measured the levels of heteronuclear, or nascent RNA as a surrogate for newly synthesized *Acsl1* transcripts indicative of transcriptional initiation [12]. We found that the Increase in steady state *Acsl1* mRNA was associated with a corresponding increase in the nascent RNA levels under HG as compared to NG conditions (Fig. 1C). This indicates that hyperglycemia induces the transcription of *Acsl1* gene in mouse BMDMs.

**Figure 1.**
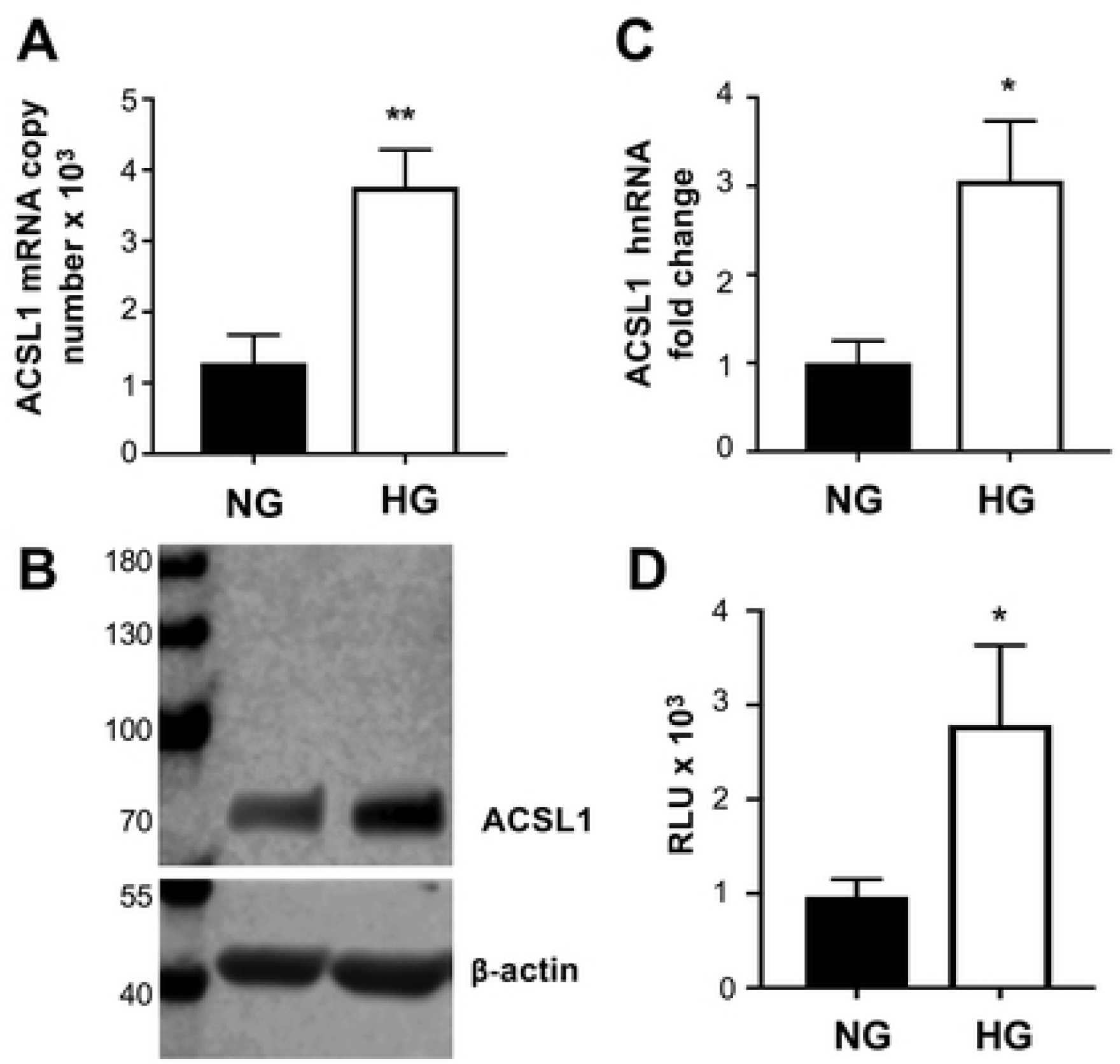
ACSL1 expression is transcriptionally upregulated in macrophages under hyperglycemic. A) BMDMs were differentiated under normal glucose (NG) and high glucose (HG) and *Acsl1* mRNA copy number was determined by quantitative real-time PCR (qPCR). B) Western blot of total cell lysates from BMDMs cultured in NG and HG using antibodies against ACSL1 and β-actin as a loading control. C) *Acsl1*nascent RNA expression was determined by qPCR using primers spanning the intron-exon junction relative to cyclophilin A1 and shown as fold change between NG and HG. D) pACSL1-GLuc reporter was transfected in HEK 293 cells cultured under NG and HG conditions. Luciferase assay was performed 48 hours post transfection, and presented a relative luciferase units (RLU). The data presented are means ± standard errors of the means of three independent experiments; the *P* value was calculated using students *t* test. Levels of significance denoted as *p < 0.05 and **p < 0.01.

To further extend these observations, we measured *Acsl1* promoter activity by performing luciferase assays in HEK293 cells transfected with an *Acsl1* construct containing the *Acsl1* promoter, and ~1.5 Kb of upstream regulatory DNA fused to the Gaussia luciferase gene (pACSL1-GLuc) under NG and HG conditions. Cells cultured in NG or HG were transfected with pACSL1-GLuc construct or with a control empty luciferase vector without the ACSL1 sequences for 48 hours before luciferase was measured. Luciferase activity was higher in cells cultured in HG as compared to NG containing the pACSL1-GLuc construct (Fig. 1D). The empty vector showed no difference in activity between high and low glucose (not shown). This is consistent with the effect of hyperglycemia on *Acsl1* mRNA expression controlled at the level of transcriptional initiation of the *Acsl1* promoter.

### CHREBP regulates *Acsl1* transcription under hyperglycemia

CHREBP is a glucose responsive transcription factor that regulates metabolic genes, including those involved in lipolysis and glycolysis [13–15]. An increase in intracellular glucose levels relieves inhibition of CHREBP and promotes CHREBP translocation from the cytoplasm into the nucleus, where it drives the expression of glucose responsive genes [16]. Elevated glucose levels in diabetes is has been shown to increase CHREBP transcriptional activity in liver and adipose tissue [17]. In addition, a ChIP-seq study for CHREBP from white adipose tissue from the fasted to fed state showed CHREBP occupies multiple sites upstream of the *Acsl1* transcription start site [18], suggesting that a *Acsl1* is a potential target of CHREBP. Because the induction of *Acsl1* by hyperglycemic is at the transcriptional level, and that CHREBP occupies an upstream regulatory region of *Acsl1* in adipose tissue, we hypothesized that the induction of *Acsl1* by hyperglycemia in macrophages is through CHREBP. To investigate this we used both gain and loss of function approaches. Prior to embarking on these experiments, we first determined the localization of CHREBP in BMDMs cultured under NG and HG conditions. We observed an increase in CHREBP nuclear localization under HG compared to NG conditions by cell fraction and immunofluorescence (S1_fig.pdf). This is consistent with CHREBP being a potential transcriptional activator of *Acsl1* under HG conditions.

Next, we used a gain of function approach to determine whether overexpression of CHREBP regulates *Acsl1* promoter activity in a cell based reporter assay. We co-transfected HEK293 cells with the same pACSL1-GLuc reporter as in Figure 1D, along with a CHREBP expression construct, or an empty expression vector, under NG and HG conditions. *Acsl1* promoter activity was higher in the cells expressing CHREBP in both NG and HG conditions (Fig 2A). The basal promoter activity in cells with vector only was also higher in the HG condition as compared to NG, and the *Acsl1* promoter activity was further increased in cells cultured in HG and overexpressing CHREBP (Fig. 2A). This is consistent with CHREBP inducing *Acsl1* transcription in HG.

**Figure 2.**
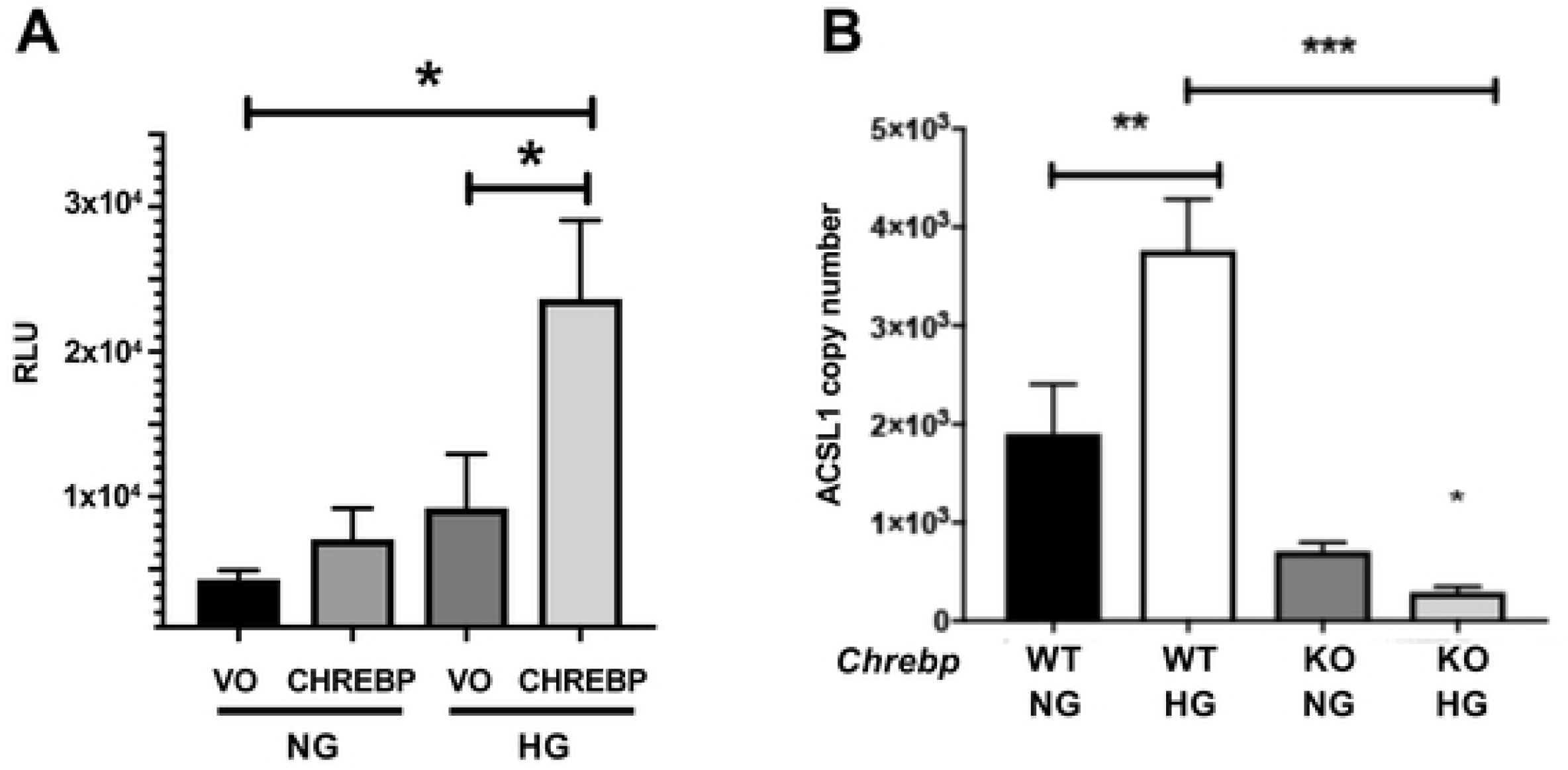
CHREBP contributes to transcriptional upregulation of ACSL1 under hyperglycemia. A) CHREP expression plasmid or vector only (VO) were co-transfected with pACSL1-GLuc reporter in HEK 293 cells cultured under NG and HG conditions. Luciferase assay was performed 48 hours post transfection and shown as relative luciferase units (RLU),. B) Wild type or *Chrebp* ^***−/−***^ BMDMs were differentiated under NG and high glucose HG conditions. *Acsl1* mRNA copy number was determined by quantitative real-time PCR (qPCR). The data presented are means ± standard errors of the means of three independent experiments; the *P* value was calculated using one way ANOVA. Levels of significance:*p < 0.05; **p < 0.01; and ***p < 0.001.

To further examine the impact of CHREBP on glucose-dependent *Acsl1* transcriptional activation we turned to a loss of function approach. We evaluated *Acsl1* expression in the absence of CHREBP under NG and HG conditions using macrophages from *Chrebp*^***−/−***^ mice [15]. BMDMs from wild type littermate and *Chrebp*^***−/−***^ mice were differentiated under NG and HG conditions, and *Acsl1* mRNA expression was measured. *Acsl1* expression was reduced in *Chrebp*^***−/−***^ cells in both NG and HG conditions, with a greater reduction in *Acsl1* expression in HG from *Chrebp*^***−/−***^ cells compared to wild type controls (Fig. 2B). This result indicates that CHREBP is required both for basal and glucose-induced expression of *Acsl1*.

### Lipopolysaccharide (LPS) stimulates *Acsl1* expression under NG and HG

Previous reports indicate that *Acsl1* expression is induced by inflammatory mediators, including LPS [7], and also is important in the TNFα-mediated inflammatory response in monocytes and macrophages [19]. It has also been reported that diabetic subjects have higher levels of *ACSL1* mRNA in circulating inflammatory monocytes [4]. Moreover, in a preclinical mouse model, myeloid specific deletion of *Acsl1* decreased the expression of proinflammatory cytokines under diabetic condition [4]. While it is evident that several pathways are capable of upregulating *Acsl1* mRNA in macrophages [3], the specific transcription factors mediating the induction of *Acsl1* via inflammatory stimuli, and how inflammatory and hyperglycemic signals intersect to promote *Acsl1* expression is not understood.

To address the regulation of *Acsl1* by inflammatory signals, we performed a time course experiment to determine the kinetics of *Acsl1* induction to LPS in macrophages. We found an induction of *Acsl1* in BMDMs between 3 and 6 hours post LPS treatment, with longer time points resulting in higher levels of *Acsl1* mRNA (S2_fig.pdf). Since a significant increase in *Acsl1* mRNA expression was observed at 24 hours, this timepoint was selected for subsequent experiments.

Given our interest in diabetes and the induction of *Acsl1* mRNA by hyperglycemia, we evaluated the impact of not only inflammation, but the combination of inflammation with hyperglycemia to *Acsl1* expression in macrophages. The results indicate that LPS treatment of macrophages (which results in M1 activated macrophages) in NG induced *Acsl1* expression ~40 fold as compared to BMDMs not activated by LPS (M0 macrophages) (Fig. 3A). Intriguingly, *Acsl1* expression upon LPS treatment in HG-induced cells increased ~80-fold relative to BMDMs not activated by LPS in NG. We also observed an increase ACSL1 protein abundance and localization to membranes upon LPS treatment (S3_fig.pdf) as has been described by others [4]. This suggests that both inflammatory and hyperglycemic signals contribute to the regulation of *Acsl1*.

**Figure 3.**
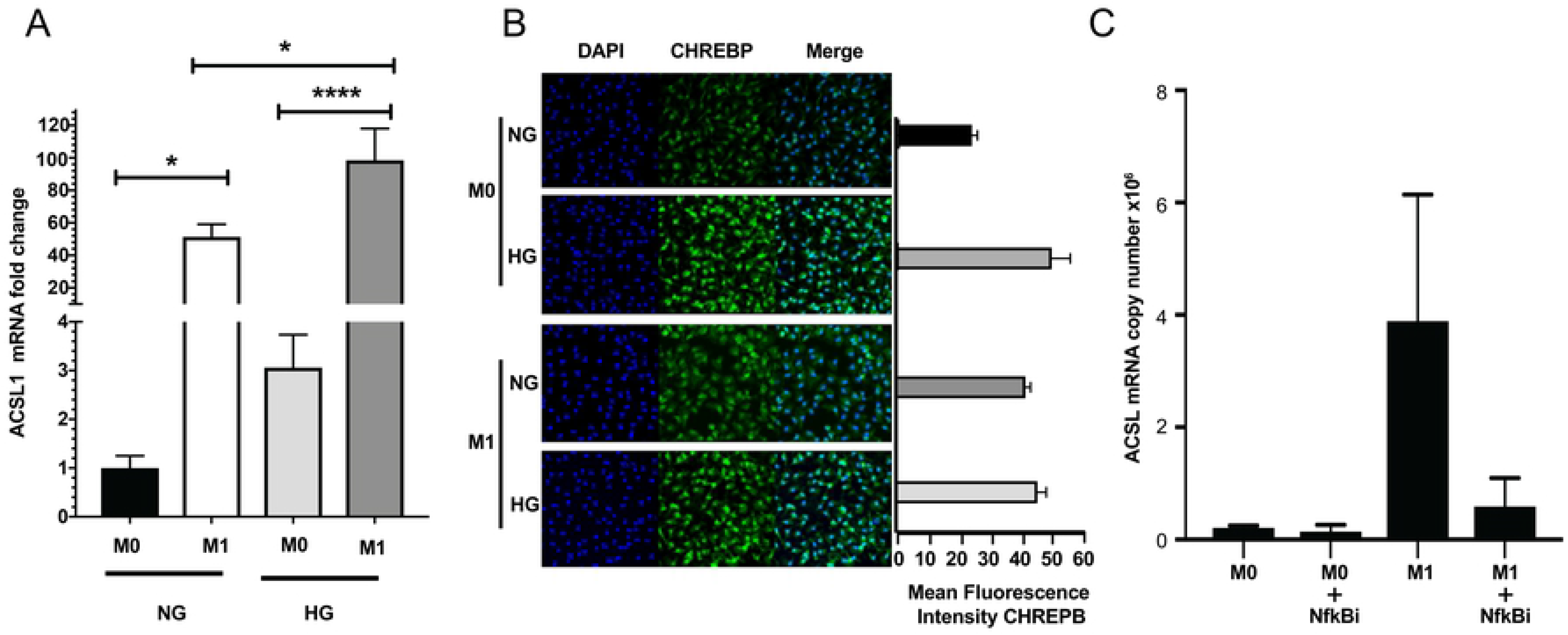
LPS induction of *Acsl1* expression is via NF-kappa B. A) BMDMs were differentiated under NG and HG conditions. Cells were treated with LPS (50ng/ml) for 24 hours. Total RNA was isolated, cDNA was synthesized and *Acsl1* mRNA was determined by qPCR relative to cyclophilin A1 mRNA and shown as fold change with M0, NG treated sample set to 1. B) BMDMs were differentiated HG conditions. Cells were pretreated for 4 hours with NF-kappa B inhibitor CAPE (5μM) and then treated with LPS at the concentration above for 16 hours. RNA was isolated, cDNA was synthesized and *Acsl1* mRNA copy number was determined by qPCR. The data presented are means ± standard errors of the means of three independent experiments; the P value was calculated using one way ANOVA. Levels of significance:*p < 0.05; **p < 0.01; and ***p < 0.001.

We also compared the localization of CHREBP protein in unstimulated (M0) and LPS-treated inflammatory (M1) macrophages under NG and HG conditions. M1 macrophages show increased nuclear CHREBP both in NG and HG conditions, with a slight increase in nuclear CHREBP under HG conditions (Fig. 3B). This suggests that CHREBP is contributing to the transcriptional regulation of ACSL1 during inflammation especially under HG conditions.

Since LPS stimulates viaToll like receptors (TLRs) the activation of NF-kappa B [20, 21], we examined whether the LPS-dependent induction of *Acsl1* was reduced by inhibition of NFκB using caffeic acid phenethyl ester (CAPE), an inhibitor that blocks NF-kappa B binding to DNA [22]. BMDMs cultured under HG conditions were either left unactivated (M0) or activated with LPS (M1) in the absence and presence of CAPE. CAPE treatment reduced LPS-dependent induction of *Acsl1* expression in macrophages (M1), while inhibiting NF-kappa B in the absence of LPS treatment did not affect the expression of *Acsl1* in M0 macrophages (Fig. 3B). This suggests that NF-kappa B activation contributes to *Acsl1* expression as a function of inflammatory stimuli, although additional factors, including but not limited to CHREBP may also participate in *Acsl1* transcription under inflammatory and hyperglycemic conditions.

### CHREBP and RELA synergistically activate*Acsl1* promoter

Given that both NF-kappa B and CHREBP appear to modulate *Acs1* expression, we examined the 1.6 kb upstream regulatory region of the mouse and human *Ascl1* genes for putative ChoRE (binding site for CHREBP) and RELA (binding site for NF-kappa B) sites using the Eukaryotic Promoter Database [8] that incorporates the Jaspar database to predict transcription factor binding site [23]. Multiple CHREBP and RELA binding sites were identified (p<0.01) in close proximity to one another that were conserved between the mouse and human genes (S4_fig.pdf). Such conservation is suggestive of the importance of the sites in transcriptional regulation [24] ,

To test the functional significance of NF-kappa B and CHREBP in establishing *Acsl1* gene expression, we employed a cell based reporter assay using the pACSL1-GLuc reporter gene containing the ~1.6 kB of upstream regulatory sequence. We transfected HEK293 cells with pACSL1-GLuc, with expression vectors for CHREBP or RELA separately, or CHREBP and RELA together, and measured the A*csl1* promoter activity. While there was a modest increase in pACSL1-GLuc activity in cells transfected with CHREBP, there was no increase in reporter activity with RELA alone (Fig. 4). Strikingly, co-expression of CHREBP and RELA showed a synergistic increase A*csl1* promoter activity (Fig. 4). This suggests that CHREBP and NF-kappa B act together to increase Acsl1 transcriptional activity.

**Figure 4.**
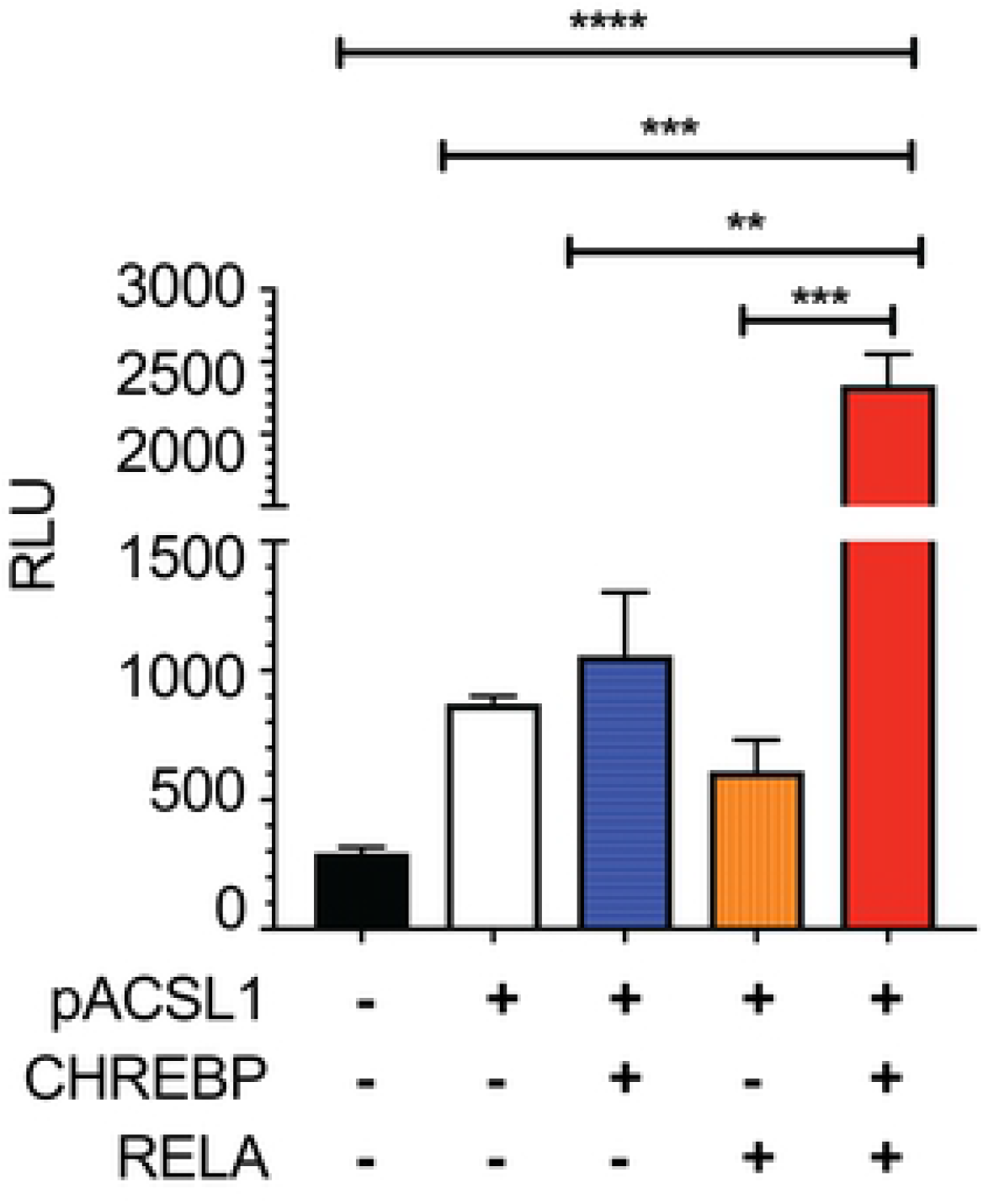
CHREBP and NF-kappa B increases ACSL1 transcriptional activity. Expression vectors of CHREBP, RELA (NFκB) or an empty vector were co-transfected with pACSL1-GLuc reporter in HEK 293 cells cultured HG conditions. Luciferase assay was performed 48 hours post transfection and shown as relative luciferase units (RLU). The data presented are means ± standard errors of the means of three independent experiments; the P value was calculated using one way ANOVA. Levels of significance:*p < 0.05; **p < 0.01; and ***p < 0.001.

## DISCUSSION

ACSL1 is one of a family of enzymes that promotes the thioesterification of long-chain fatty acids to form acyl-CoAs for use in synthetic or degradative pathways [2, 3]. In metabolically active tissues the acyl-CoAs go toward mitochondrial ß-oxidation. ACSL1 can also specify phospholipid synthesis, and these can serve as a source of arachidonic acid-CoA metabolites that support prostaglandin synthesis to fuel inflammation in diabetes that exacerbates atherosclerosis [4]. Whereas *Acsl1* mRNA expression is controlled by inflammation and hyperglycemia, the mechanisms underlying the induction of *Acsl1* mRNA by these stimuli have remained enigmatic. In this study, we defined transcription factors controlling *Acsl1*gene expression in macrophages. We show that *Acsl1*expression is regulated by CHREBP in hyperglycemia and through NF-kappa B under inflammatory conditions. This is consistent with recent reports showing increased ACSL1 mRNA in peripheral blood from septic patients [25], and also elevation in blood of patients with acute myocardial infarction compared to controls [11], which may reflect an inflammatory response from necrotic tissue as a result of ischemia.

CHREBP appears to control *Acsl1* mRNA expression in both NG and HG in M0 macrophages, suggesting even under NG there is some active CHREBP, which becomes increased in HG to promote *Acsl1* transcription. This suggests that CHREBP under hyperglycemia is a key determinant in the increased expression of *Acsl1* observed in monocytes and macrophages from both humans and mice under conditions of diabetes. This is reflected in the glucose-dependent induction of *Acsl1* steady state mRNA and nascent RNA expression, and the overexpression and deletion of CHREBP that increased and decreased, respectively, *Acs1* transcriptional activity.

Acute inflammatory stimuli by LPS in macrophages promotes a robust induction of *Acsl1* mRNA. This can be largely suppressed by an NF-kappa B inhibitor. Although it appears that induction of *Acsl1* transcription by LPS predominates relative to the expression under hyperglycemia, the inflammatory response of *Acsl1* expression can be further enhanced in hyperglycemic conditions, suggesting an interplay between pathways. Consistent with this is the synergistic increase in *Acsl1* promoter activity when both CHREBP and RELA were co-expressed, suggesting the two factors cooperate to drive *Acs1* gene expression in settings of inflammation and hyperglycemia. This is also in line with binding sites for both factors being present in the upstream ACSL1 regulatory region and conserved between the mouse and human genes. Such conservation is often sufficient to predict transcription factor occupancy and activity at induced genes [24]. Indeed, CHREBP has been shown to occupy the *Acsl1* upstream regulatory region in mouse adipose tissue [18], and analysis of the human ACSL1 gene from ENCODE shows RELA occupancy in the region upstream of the ACSL1 start site of transcription in a variety of cells types. This is reinforced by the functional studies that indicate both CHREBP and NF-kappa B enhance the expression of *Acsl1*.

Based on these findings, we propose that in macrophages under conditions of NG and either no or low level inflammation the expression of *Acsl1* is low and driven by a small pool of active, nuclear CHREBP (Fig. 5A). Expression of *Acsl1* is increased under conditions of HG by virtue of an increase in the pool of nuclear CHREBP (Fig. 5B). We further posit that in macrophages exposed to an acute inflammatory stimuli, such as by LPS, expression is of *Acsl1* is increased by activation of NF-kappa B under both NG and HG, with even greater activation in HG (Fig. 5C). Whether NF-kappa B induction of *Acsl1* also requires CHREBP remains an open question, but is suggested by the low activity of the *Acsl1* reporter with overexpressed RELA, and that co-expression of both CHREBP and RELA synergistically activate the *Acsl1* reporter. Thus, our studies have revealed the convergence of two important pathways on the regulation of *Acsl1* in macrophages to align the production of acyl-CoA derivatives to the cellular environment.

**Figure 5.**
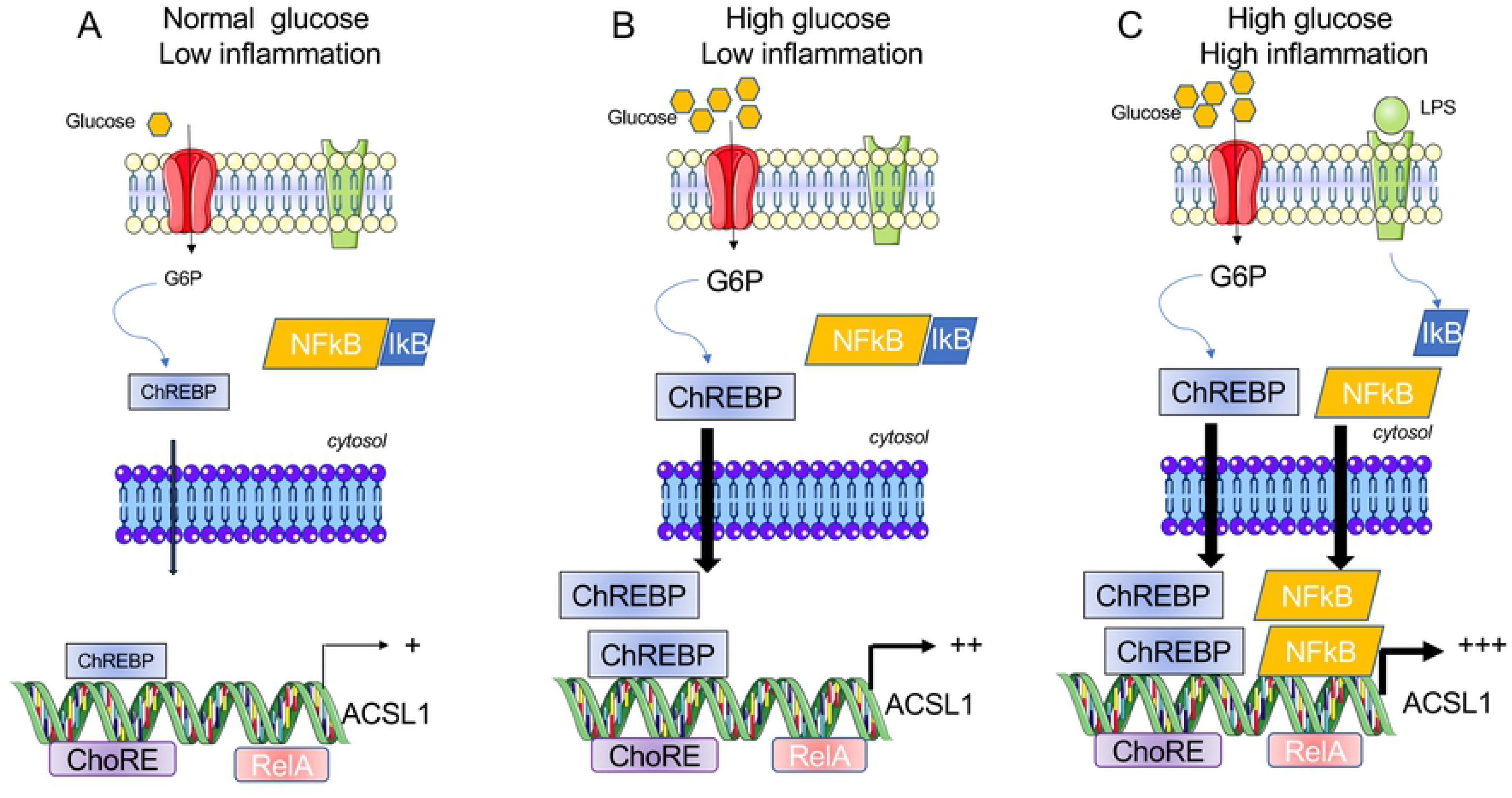
Model for glucose and inflammation induced *Acsl1* expression by CHREBP and NFκB in macrophages. We propose that under A) normal glucose and low inflammation, there is a small amount active CHREBP in the nucleus that promotes the expression of *Acsl1*. B) In high glucose this pool increases to promote a higher level of *Acsl1* mRNA. C) During acute inflammation, such as with treatment with LPS, NF-kappa B, which is normally held in check by Inhibitor of kappa B (ikB), is free to translocate to the nucleus where it activates transcription of *Acsl1,* which can be accentuated by CHREBP.

## Author contributions

Prashanth Thevkar-Nagesh; was involved with the investigation and analysis of the data, as well as writing the original draft preparation of the manuscript; Shruti Rawal and Tarik Zahr were involved in the investigation; Edward A. Fisher and Michael J. Garabedian were involved with conceptualization, funding acquisition, project administration, supervision, writing the original draft as well as its review and editing the final version of the manuscript.

## Acknowledgements

We thank Drs. Claudia Han and Christopher Glass (UCSD) for providing the CHREBP knock out macrophages. We also thank Hussam Alkaissi (NYU) for help with the ACSL1 membrane localization experiment. This work was supported by an NIH grant (P01HL131481) to E.A.F and M.J.G.

**S1. Increased nuclear localization of CHREBP in macrophages in hyperglycemia.** A) BMDMs were differentiated under NG and HG conditions. Cytoplasmic and membrane proteins were isolated and western blot performed using an ant-CHREBP to determine the abundance and subcellular localization of CHREBP protein. Tubulin and Histone H3 serve as controls for cytoplasmic and nuclear fractions. B) BMDMs were differentiated under NG and HG conditions. The cells were grown on cover slips and stained for CHREBP and DAPI to visualize the nucleus. The images were obtained using Leica SP5 Confocal Microscope at 63X magnification.

**S2. Kinetics of ACSL1 induction by LPS treatment.** BMDMs were differentiated in NG and treated with LPS (50ng/ml). RNA was isolated at the indicated times, cDNA was synthesized and *Acsl1* mRNA expression was determined by qPCR. The data presented are means ± standard errors of the means of two independent experiments. Levels of significance:*p < 0.05; **p < 0.01; and ***p < 0.001.

**S3. ACSL1 protein abundance and membrane localized increase under inflammatory conditions.** BMDMs were differentiated under NG conditions. A)Total cell lysates were prepared after LPS treatment (50ng/ml for 24 hours) and ACSL1 protein abundance was determine by western blot with an anti-ACSL1 antibody and with an anti-tubulin antibody as a loading control. B) Bands were quantitated using the Image studio 5.2 and plotted using Graph pad prism. The data presented are means ± standard errors of the means of three independent experiments. C) Cytoplasmic and membrane proteins were isolated to determine the localization of ACSL1 protein by Western blotting upon LPS treatment. D) Image studio 5.2 was used to quantify the bands and the values were plotted using Graph pad prism. The data presented are means ± standard errors of the means of three independent experiments.

**S4. LPS treatment increases CHREBP abundance in the nucleus.** BMDMs were differentiated under NG and HG conditions. Cells were grown on cover slips, treated with LPS (50ng/ml) for 24 hours and stained for CHREBP. The images were obtained using Leica SP5 Confocal Microscope at 63X magnification. Image J was used to quantify the fluorescence intensity and Graph Pad Prism was used to plot the fluorescence intensities.

**S5. Predicted CHREBP and NFκB sites upstream of ACSL1 promoter in the human and mouse genes.** The Eukaryotic Promoter Database tool was used to predict the putative CHREBP and NFκB sites in human and mice upstream ACSL1 promoter region via the Jaspar database. The P-value used in the prediction was p<0.01.

